# The genetic diversity of the soybean rust pathogen *Phakopsora pachyrhizi* has been driven by two major evolutionary lineages

**DOI:** 10.1101/2024.10.03.616435

**Authors:** Vinicius Delgado da Rocha, Everton Geraldo Capote Ferreira, Fernanda Machado Castanho, Marcia Kamogae Kuwahara, Cláudia Vieira Godoy, Maurício Conrado Meyer, Kerry F. Pedley, Ralf T. Voegele, Anna Lipzen, Kerrie Barry, Igor V. Grigoriev, Marco Loehrer, Ulrich Schaffrath, Catherine Sirven, Sebastien Duplessis, Francismar Corrêa Marcelino-Guimarães

**Affiliations:** Brazilian Agricultural Research Corporation - National Soybean Research Center (Embrapa Soja), Paraná, Brazil; Blades, Evanston, Illinois, USA; The Sainsbury Laboratory, University of East Anglia, Norwich, UK; United States Department of Agriculture-Agricultural Research Service (USDA-ARS), Foreign Disease-Weed Science Research Unit, Ft. Detrick, Maryland, U.S.A; Department of Phytopathology, Institute of Phytomedicine, Faculty of Agricultural Sciences, University of Hohenheim, 70599 Stuttgart, Germany; United States Department of Energy Joint Genome Institute, Lawrence Berkeley National Laboratory, Berkeley, California, U.S.A; Department of Plant and Microbial Biology, University of California Berkeley, Berkeley, California, U.S.A; Department of Plant Physiology, RWTH Aachen University, Aachen, Germany; Bayer SAS, Crop Science Division, Lyon, France; Université de Lorraine, INRAE, IAM, Nancy, F-54000, France

**Keywords:** genetic admixture, mating-type genes, plant-pathogens, population genomics, virulence

## Abstract

*Phakopsora pachyrhizi*, an obligate biotrophic rust fungus, is the causal agent of Asian Soybean Rust (ASR) disease. Here, we utilized whole-genome data to explore the evolutionary patterns and population structure across 45 *P. pachyrhizi* isolates collected from 1972 to 2017 from diverse geographic regions worldwide. We also characterized *in-silico* mating-type (*MAT*) genes of *P. pachyrhizi*, in the predicted proteome of three isolates, to investigate the sexual compatibility system. Our molecular phylogenetic analysis in *P. pachyrhizi* inferred two distinct evolutionary lineages structured on a temporal scale, with lineage Pp1 grouping isolates obtained from 1972 to 1994, while more recently collected isolates formed a second lineage, Pp2. We found high levels of genetic diversity in lineage Pp1, whereas lineage Pp2 exhibited a strong clonal genetic structure, with a significant lower diversity. The widespread propagation of *P. pachyrhizi* clonal spores across soybean-growing regions likely explains the absence of a large-scale spatial genetic structure within each lineage. Two independent isolates (TW72-1 and AU79-1) showed moderate levels of genetic admixture, suggesting potential somatic hybridization between the two *P. pachyrhizi* lineages. We observed no clear congruence between virulence levels of *P. pachyrhizi* isolates and their phylogenetic patterns. Our findings support a probable tetrapolar mating system in *P. pachyrhizi*. Taken together, our study offers new insights into the evolutionary history of *P. pachyrhizi* and demonstrates that multiple *MAT* genes are highly expressed during the later stages of soybean infection, suggesting their potential role in the formation of urediniospores within the life cycle of *P. pachyrhizi*.

**AUTHOR SUMMARY:** The Asian Soybean Rust (ASR) disease, caused by basidiomycetes fungus *P. pachyrhizi*, represents a critical threat to soybean crops worldwide. With the recent availability of high-quality genome assemblies for *P. pachyrhizi*, we are committed to exploring the genetic diversity of this destructive pathogen. This study analyzed whole-genome resequencing data from *P. pachyrhizi* isolates collected over several decades in various geographic regions. We identified recent diversification patterns in *P. pachyrhizi*, with two major lineages. The origin of the two lineages is likely due to temporal shifts in the genetic structure of *P. pachyrhizi*. Additionally, we investigated the genes responsible for sexual compatibility, known as mating-type (*MAT*) genes. Transcriptome data indicated that *MAT* genes are actively expressed during the later stages of the *P. pachyrhizi*-soybean infection. We conclude that the evolutionary dynamic of *P. pachyrhizi* is shaped by divergent lineages, which exhibit varying levels of virulence on differential soybean lines containing ASR resistance genes.

## INTRODUCTION

The rust fungus *Phakopsora pachyrhizi* (Phakopsoraceae) is an obligate biotrophic plant-pathogen known for its ability to infect around 150 plant species within the family Fabaceae, including soybean (*Glycine max*) [1]. The broad host range of *P. pachyrhizi* may play a crucial role in its successful adaptability, serving as an inoculum reservoir for survival and dissemination of the pathogen on other crops and between planting seasons when the primary host (i.e. soybean) is not available [2]. Notably, *P. pachyrhizi* is the causal agent of Asian Soybean Rust (ASR), one of the most devastating foliar diseases to soybean production, especially in South America, where favorable environmental conditions contribute to the development of the disease. Without adequate management, ASR can lead to severe yield losses, reaching up to 80% [3,4]. ASR symptoms consist of grayish-brown lesions that form predominantly on the abaxial surface of leaves, prompting intense premature leaf senescence [5]. The control and management of ASR is particularly challenging due to the high variability in virulence levels of *P. pachyrhizi*, with the potential occurrence of multiple races [6, 7]. Although seven loci (*Rpp1*-*Rpp7*) conferring resistance to ASR have been described in soybeans, *P. pachyrhizi* has rapidly evolved to overcome individual resistance genes [8].

*Phakopsora pachyrhizi* likely originated on the Asian continent and subsequently spread to other worldwide soybean-growing regions, including Africa and the Americas [5]. The first documented occurrence of the pathogen dates back to 1902 in Japan, where it was initially identified as *Uredo sojae* [9]. Since then, the presence of *P. pachyrhizi* has been observed in several geographically distinct regions. The pathogen was reported in India in 1970 [10], in Africa during the 1990s [11, 12], South America in 2001 [13], and North America in 2004 [14].

The global dispersion of *P. pachyrhizi* has been associated with intercontinental dispersals of airborne spores [14, 15]. Indeed, evidence-based studies suggest that transatlantic air currents may have promoted the widespread dissemination of *P. pachyrhizi* spores from Africa to Brazil [16]. The occurrence of long-distance dispersal events for *P. pachyrhizi* has shaped its patterns of genetic diversity and population structure, leading to intense gene flow among spatially separated populations [16, 15, 17]. Previous investigations based on molecular markers, such as microsatellites [17], AFLPs (Amplified Fragment Length Polymorphism) [18], ITS (Internal transcribed spacer), and ARF (ADP-Ribosylation Factor) gene sequences [19], have demonstrated that *P. pachyrhizi* isolates from various localities exhibited low levels of genetic differentiation across extensive geographic areas in Africa, North America, and South America, implying a predominantly clonal population structure. Similarly, comparisons among three *P. pachyrhizi* isolates from South America (K8108, MT2006, and UFV02), based on single nucleotide variations distributed across their genomes, demonstrated that they potentially represent a single clonal lineage with high levels of heterozygosity [20].

Whether sexual recombination contributes to the diversity of *P. pachyrhizi* remains an open question, as its sexual cycle is completely unknown. *P. pachyrhizi* reproduces primarily, if not exclusively, via asexual urediniospores that harbor two separate haploid nuclei (N + N) [21]. Teliospores are the only other spore type that has been identified in *P. pachyrhizi*; however, their germination has never been observed in nature [21, 22].

The sexual compatibility patterns of *P. pachyrhizi*, particularly involving mating-type (MAT) genes, remain to be elucidated. Conversely, comparative genomic analyses have identified MAT genes in other rust fungi, including *Melampsora larici-populina* [23], various *Puccinia* species [23, 24, 25], and *Austropuccinia psidii* [26]. In general, available genomes of plant-pathogenic rust fungi, (e.g. *Puccinia* species), show a tetrapolar compatibility system, with MAT genes organized into two loci — P/R and HD locus — located on distinct chromosomes [25]. Specifically, the P/R locus contains genes-encoding pheromone peptide precursors (MFA), as well as multiple genes that encode pheromone receptor proteins known as STE3. The HD locus carries alternative alleles that encode two divergent homeodomain transcription factors (HD1 and HD2) [27]. Within the Basidiomycota, the phylum in which *P. pachyrhizi* is included, sexual mating compatibility involves molecular recognition between mature pheromone peptides and their corresponding transmembrane G-protein coupled receptors [27]. This recognition triggers the fusion of compatible gametes and subsequently leads to the formation of a heterodimeric complex composed of HD1 and HD2 homeodomain transcription factors [24, 27]. The HD1-HD2 heterodimer acts as a transcriptional regulator. It binds to promoter sequences and enhances the expression of specific genes, thus promoting sexual development [28].

The first genome assemblies of *P. pachyrhizi* have been recently published [20]. By combining two different sequencing platforms — short-read data and long- read data — robust bioinformatics tools constructed *de novo* genome assemblies of three distinct isolates of *P. pachyrhizi* (K8108, MT2006, and UFV02) [20]. These genomic resources can be used to gain insight into the evolutionary landscape of the pathogen over spatial and temporal dimensions.

In this study, we elucidated the evolutionary patterns and population structure of *P. pachyrhizi*. To achieve this objective, we utilized the Illumina sequencing platform to re-sequence whole-genomes of 42 diverse isolates of *P. pachyrhizi*. In addition, we compiled genomic data for three previously investigated isolates: K8108, MT2006, and UFV02 [20]. Our collection of isolates was acquired within a wide timeframe (1972-2017) and covers a global geographic range across various soybean-growing areas, comprising Africa, Asia, Australia, South America, and the United States of America (USA). We identified highly-curated SNPs at the whole-genome level by mapping the *P. pachyrhizi* isolates to the UFV02 reference genome. Next, we implemented phylogenetic reconstructions and population genomic approaches to determine whether multiple evolutionary lineages have influenced the distribution patterns of genetic diversity among *P. pachyrhizi* isolates over the course of time. To explore whether distinct evolutionary lineages of *P. pachyrhizi* may potentially outcross through sexual compatibility, we performed *in-silico* functional annotation of MAT genes of *P. pachyrhizi,* and their expression profiles were investigated during the interaction between the pathogen and soybean host. We also carried out phenotyping experiments to assess the virulence levels of *P. pachyrhizi* isolates on differential soybean genotypes containing seven loci that confer resistance to *P. pachyrhizi* (*Rpp1 to Rpp7*). Our study enables us to expand our current understanding of how evolutionary processes have shaped the genomic diversity of *P. pachyrhizi*.

## MATERIAL AND METHODS

### Fungal isolate sampling

Our study encompassed 37 *P. pachyrhizi* isolates, which were deposited in the Collection of Microorganisms of the Embrapa Soybean (CMES), Londrina, Paraná, Brazil. The isolates were sampled over 15 years (2003-2017) and obtained from soybean leaves collected in four geographically distant regions, including Africa, Asia, South America, and the USA (S1 Table). Each isolate was subjected to single-spore purification on detached soybean leaflets. In summary, a drop of water was carefully placed over a single uredinium. Single spores were isolated via a Zeiss Axio Scope A1 microscope. A single spore was transferred to healthy leaflets of the susceptible genotype Embrapa 48, kept in a Petri dish containing 1% agar-water medium, and then maintained in a growth chamber, at a temperature of approximately 21°C and a humidity level of 60% for approximately 15 days for the production of new lesions. To guarantee the purity of the isolates, three cycles of single spore isolation were carried out. Detached leaflets of the same susceptible genotypes were used for spore multiplication, and the spores were transferred to a 2 mL microtube kept open in a desiccator for 24 hours for dehydration and were then stored in a freezer at −80°C for preservation.

Additionally, we also included five *P. pachyrhizi* isolates representing early strains that were originally collected between the years 1972 and 2004 (S1 Table). These five isolates [Taiwan-1972 (TW72-1), India-1973 (IN73-1), Australia-1979 (AU79-1), Hawaii-1994 (HW94-1), and Colombia-2004 (CO04-2)] were maintained at the United States Department of Agriculture (USDA), Foreign Disease-Weed Science Research Unit (FDWSRU), Frederick, MD, USA.

### DNA extraction and whole-genome sequencing

Total genomic DNA was extracted from 2-3 mg of spore per isolate, following an existing protocol with adaptations [29]. DNA concentration was quantified by a Nanodrop 8000 spectrophotometer (Thermo Scientific, Wilmington, DE). Whole-genome sequencing of each isolate was performed at DOE Joint Genome Institute (JGI), using paired-end sequencing (2 x 151 bp) on an Illumina NovaSeq S4 platform. To evaluate the quality of raw sequence data, we employed a JGI QC pipeline to remove potential contaminants, adapters, and low-quality sequences.

### Mapping and variant calling

Both read-mapping and variant detection were conducted according to the procedures developed by a previous study [20]. Illumina sequence data from each isolate were mapped to the *P. pachyrhiz*i UFV02 v2.1 reference genome [20] using the BWA-MEM algorithm with options -M-R in BWA v. 0.7.17 software [30]. The SAMtools v1.9 software [31] was employed to convert alignment files into compressed BAM format. Subsequently, bam files were input into the Picard package (https://broadinstitute.github.io/picard/) to remove duplicate reads. Local realignments of sequencing-reads around indel regions were performed using two algorithms (RealignerTargetCreator and IndelRealigner) implemented in the GATK v3.8.1 software [32].

SNPs and indels were identified using the HaplotypeCaller algorithm in GATK software, producing a single VCF file for 42 *P. pachyrhiz*i isolates. To ensure high-quality variants, the resulting VCF file was filtered in GATK v.4.2 software [32], with the following thresholds: QUAL < 30.00, MQ < 40.00, SOR > 3.00, QD < 2.00, FS > 60.00, MQRankSum < -12.500, ReadPosRankSum < -8.00. Utilizing VCFtools v0.1.16 software [33], the VCF file underwent additional filtering parameters to remove indels and retain biallelic SNPs with a minimum depth of read coverage (minDP = 5) and minimum allele frequency (MAF ≥ 5%).

To incorporate additional genomic data into our study, we generated a SNP set based on publicly available genomic assemblies of three *P. pachyrhizi* isolates, K8108, MT2006, and UFV02 [20]. This analysis utilized the NUCmer algorithm implemented in MUMmer v3.23 [34], with options –show-snps -Clr. Using the “merge” function in the BCFtools software [35], the predicted SNPs in the three isolates were concatenated with our VCF file, comprising the SNPs from 42 isolates. Finally, we obtained a large SNP panel containing 452,853 whole-genomic variants (MAF ≥ 5% and missing data ≤ 20%) across 45 *P. pachyrhizi* isolates. We referred to this SNP panel as Dataset A.

### Annotation and prediction of SNP effects

We applied SnpEff v.5.1 software [36] to annotate and predict the putative effects of SNPs on coding-DNA regions in *P. pachyrhiz*i. The UFV02 v2.1 reference genome and its gene annotation GFF file [20] were utilized to build a SnpEff database through the “built” function and default parameters. Dataset A served as an input file for SnpEff.

### Phylogenetic analyses and population structure

Maximum likelihood (ML) approaches reconstructed the phylogenetic patterns among 45 *P. pachyrhizi* isolates by analyzing Dataset A. Before achieving phylogenetic analyses, the best-fit model of nucleotide evolution was inferred through the ModelFinder2 software tool [37] implemented in IQ-TREE v1.6.12 [38]. The Bayesian Information Criterion (BIC) chose TVMe as the best-fit model. We conducted an ML phylogeny in IQ-TREE software, with ten independent runs and 1,000 ultrafast bootstrap replicates. The bootstrap convergence criterion was 0.99 and followed the standards described by a previous work [39]. The resulting phylogenetic tree was edited using FigTree v1.2.4 (http://tree.bio.ed.ac.uk/software/figtree/).

To reduce linkage disequilibrium (LD) bias on the population structure analyses, we selected randomly SNPs separated within a distance of 5 kbp utilizing parameter --thin 5000 in VCFtools v0.1.16 [33]. The SNPs were filtered with MAF ≥ 5% and missing data ≤ 20%, generating a final VCF file containing 115,065 whole-genomic SNPs. From this data, we assembled two distinct datasets — Dataset B and Dataset C — to explore population structure in *P. pachyrhizi*. Dataset B came from a full set of 45 isolates, while Dataset C included exclusively 41 isolates belonging to lineage Pp2 uncovered during the preceding phylogenetic analysis (Fig 1A).

**Fig 1.**
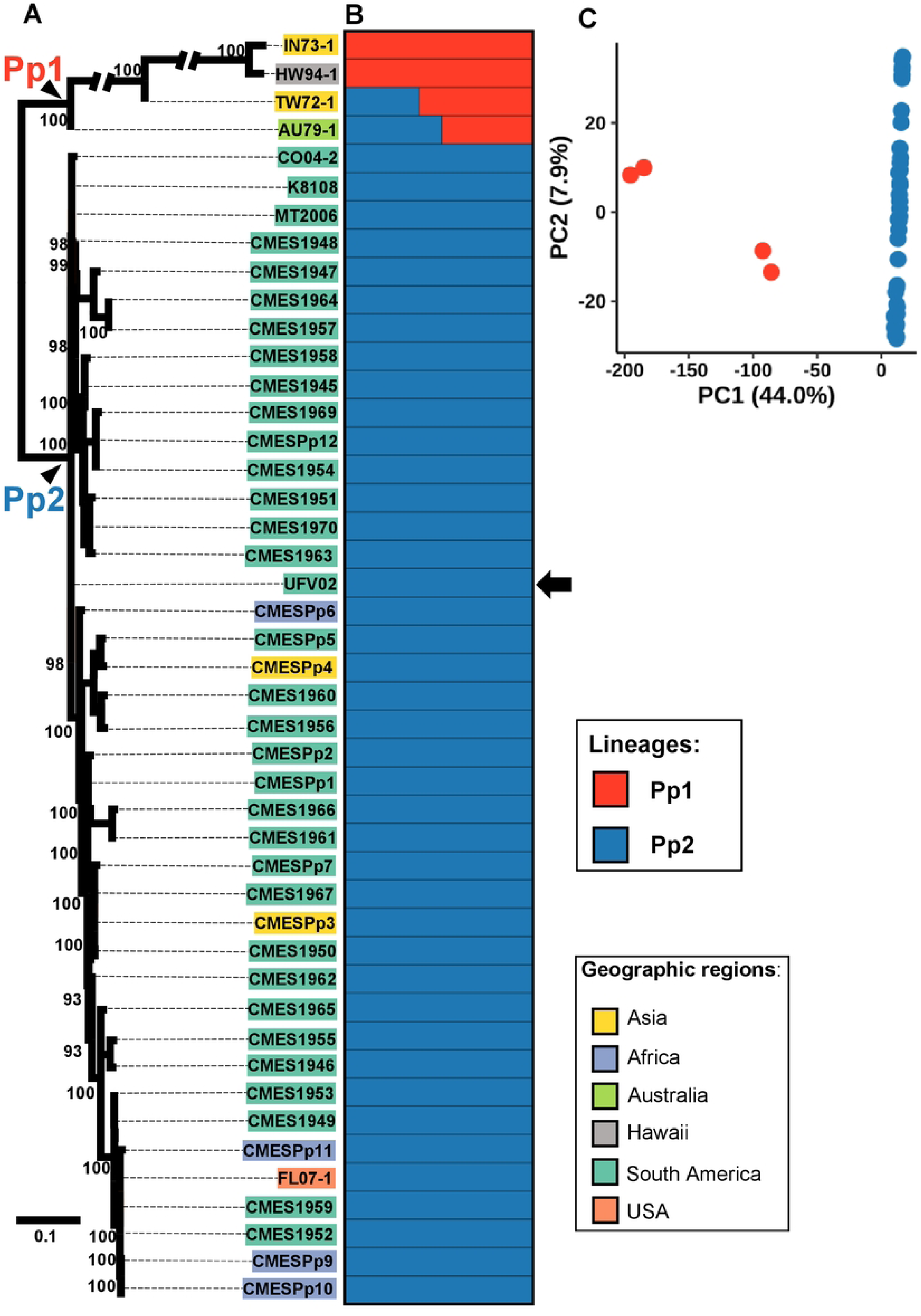
Phylogenetic reconstruction and population structure in *Phakopsora pachyrhizi.* The analyses utilized whole-genome SNP data from 45 isolates collected from six geographic regions (Asia, Africa, Australia, Hawaii, South America, and the USA). **A.** The maximum likelihood tree (unrooted) reveals two major clades, designated as lineage Pp1 and lineage Pp2. Branch lengths are proportional to the scale bar that corresponds to the expected number of substitutions per site. Bootstrap support values are displayed above the branches (when > 80). The arrow indicates the phylogenetic placement of the UFV02 reference genome. Isolate names are color-coded according to geographic regions, as indicated. **B.** fastSTRUCTURE analysis points out the presence of two Bayesian clusters, as supported by optimal K number (K=2). Each colored horizontal bar represents the ancestry proportion of an isolate to its respective Bayesian cluster. **C.** Plot of the first two principal components, with eigenvalues for each component shown in parentheses.

Population structure analyses were conducted with two independent methods: principal component analysis (PCA) and Bayesian clustering. PCA was performed using R v4.4 (https://www.r-project.org/), with the “glPca” function implemented in the adegenet package v2.1.10 [40]. The 2D PCA plot was generated in the R ggplot package v3.5.1 [41]. The fastStructure v1.0 [42] was utilized to determine Bayesian model-based clustering, ranging K from 1 to 10, with 10 independent replications for each K. The optimal number of K was chosen based on a FastStructure chooseK.py script. The Q-matrix files from 10 replications, representing membership probabilities for the optimal K, were merged and results were visualized in the R pophelper package v2.3.1 [43].

### Genetic diversity analyses

To calculate the pairwise genetic differentiation index (F_ST_) between two *P. pachyrhizi* lineages identified in our initial phylogenetic inference (Fig 1A), we carefully divided Dataset A into two subsets, Dataset D and Dataset E, each harboring 307,752 whole-genomic SNPs from distinct lineage P1 and P2, respectively. For these datasets, we applied a new filter with MAF ≥ 5% and sites with missing data were discarded. The pairwise FST values [44] were estimated in VCFtools v0.1.16 [33], with non-overlapping sliding windows of 10 kbp.

To assess genetic diversity levels among isolates of each *P. pachyrhizi* lineage, we calculated two independent measures: nucleotide diversity per site (π) and Watterson’s theta per site (θw). We also computed the Tajima’s D test neutrality to detect potential signals of selective pressure acting on *P. pachyrhizi* lineages. These analyses were conducted in the R PopGenome package v2.7.5 [45] and were structured into two stages. First stage, we included Dataset D and Dataset E containing SNPs distributed across the whole-genome. Second stage, we included exclusively SNPs located in genes (both predicted effector and non-effector), utilizing the annotated GFF3 file of the UFV02 isolate as a reference gene annotation [20]. The classification of candidate effector genes in the UFV02 isolate was based on its predicted secretome, with well-established criteria [see 20]. Specifically, we considered a total of 1,168 candidate effector genes predicted in the UFV02 isolate, while 21, 299 remaining genes were categorized as non-effectors [20].

The non-parametric Mann-Whitney-Wilcoxon test inferred statistical differences between the two *P. pachyrhizi* lineages for genetic diversity measures and Tajima’s D values, utilizing R v4.4 (https://www.r-project.org/).

### Identification of mating-type (*MAT*) genes

For *in-silico* characterization of mating-type (*MAT*) genes in *P. pachyrhizi*, we used the predicted proteomes from the three isolates K8108, MT2006, and UFV02 [20]. We ran local BLASTp searches (cutoff e-value < 1e-5) to predict putative *MAT* genes, encompassing pheromone receptors (*STE3.2*), pheromone precursors (*MFA*), and homeodomains (*HD*). As queries, BLASTp searches utilized MAT protein sequences previously identified in two closely related rust fungal species: *Puccinia triticina* and *Puccinia graminis* f. sp. *tritici* [24] (S2 Table). Furthermore, we also searched for *MAT* homologs through HMMER v3.3.2 software (http://hmmer.org/), with an e-value < 0.001, employing independent profile Hidden Markov Models for Pheromone A receptor (PF02076) and Homeobox KN domain (PF05920), downloaded from the Pfam database v33.0 [46].

After identifying putative *MAT* genes in *P. pachyrhizi*, we examined their phylogenetic relationships among the isolates K8108, MT2006, and UFV02, and two *Puccinia* species were used as outgroups. For this purpose, we created three datasets, each consisting of protein sequences from a different *MAT* gene in *P. pachyrhizi*. The MAT protein sequences were aligned utilizing MAFFT v7.453 [47] under the L-INS-i algorithm, with default parameters. For each set of *MAT* genes, independent ML phylogenies were performed in IQ-TREE v1.6.12 [38], with ten separate runs and 1,000 ultrafast bootstrap replicates.

To determine whether the MAT genes of *P. pachyrhizi* were differentially expressed throughout the soybean infection cycle, we retrieved transcriptome data for the three *P. pachyrhizi* isolates—K8108, MT2006, and UFV02—available as Log2 fold change values of gene expression during different timepoints of spore germination and soybean infection [20]. Heatmap plots were constructed using the R ggplot2 package to visualize the expression profiles of *MAT* genes in *P. pachyrhizi*.

### Virulence of *P. pachyrhizi* isolates on differential soybean genotypes

We performed comprehensive phenotyping evaluations to investigate the virulence profiles of seven *P. pachyrhizi* isolates (TW72-1, IN73-1, AU79-1, HW94-1, CO04-2, MT2006, and UFV02) from the two main evolutionary lineages recognized in our study (Fig 1A). To characterize the pathogenicity of *P. pachyrhizi* isolates, we used eight differential soybean accessions, each carrying a single *Rpp* gene. The complete list of soybean plant introduction (PI) accessions used in this study can be found in S3 Table.

Phenotyping analyses for isolates MT2006 and UFV02 were carried out at Embrapa Soybean, Londrina, Brazil. In a greenhouse, we cultivated three plants per soybean accession until they reached the V3-V4 growth stage. The third trifoliate of each plant was collected and kept in Petri dishes with 1% agar-water medium for subsequent inoculation. For each fungal isolate, we prepared a spore suspension (4 × 10^4^ spores/mL) consisting of sterile distilled water and 0.01% Tween® 20. The spore suspension was spread homogeneously on the abaxial side of leaves using a glass atomizer. The inoculated leaves were kept in a grow chamber with 60% relative humidity and 24°C for 14 days. After the incubation period, leaves were photographed and evaluated based on four previous quantitative parameters [48]. The four parameters were as follows: (1) presence or absence of lesions, (2) lesion sporulation level (SL), (3) number of uredinia per lesion (NoU), and (4) frequency of lesions showing uredinia (%LU) (S4 Table). In each soybean accession, 3 biological replicates (one leaf per plant) were analyzed, with 30 lesions per leaf, totaling 90 lesions per soybean accession. Based on the quantitative parameters, four qualitative categories were established: immune (I), highly resistant (R), intermediate resistant (IM), and susceptible (S). As a control, we used the susceptible cultivar BRS 184.

For the five remaining isolates (AU79-1, CO04-2, HW94-1, IN73-1, and TW72- 1), we conducted phenotyping assays at the FDWSRU. Soybean plants were grown in a growth chamber (Environmental Growth Chambers) at a constant temperature of 20**°**C with a 16-h photoperiod. All plants were fertilized 3 weeks after germination with Peters Professional 20-20-20. Plants were transferred to the USDA biological safety level 3 (BSL-3) plant pathogen containment greenhouse facility at Ft. Detrick and inoculated with *P. pachyrhizi* approximately 3 weeks after germination. Respective isolates were prepared as previously described [49]. Spore concentrations were measured with a hemacytometer and adjusted to approximately 5×10^4^ spores/mL in a solution of 0.01% Tween® 20 in sterile distilled water. Inoculum was sprayed onto plants with an atomizer until leaves were saturated. Inoculated plants were incubated overnight in a 20°C dew chamber, and then transferred to a 25°C greenhouse. The susceptible line Williams 82 (Wm82) was used as a control. Plants were photographed 14 days after inoculation. Phenotypes were scored as immune (IM1), immune with flecking (IMF), red-brown lesions with no uredinia (RB2), red-brown lesions with some lesions containing uredinia (RB3), red-brown lesions with most lesions containing uredinia (RB4), or Tan lesions with uredinia (TAN5). A mixed phenotype was observed for one soybean line and *P. pachyrhizi* isolate combination.

To conduct subsequent statistical analyses, we constructed a data matrix by attributing numerical scores to the phenotypic responses of soybean genotypes against seven *P. pachyrhizi* isolates. This matrix classified disease severity degree into six distinct classes, each of which was converted to a numerical score: 0 = Immune, 1 = Immune with flecking, 2 = Red-brown lesions with no uredinia (highly resistant), 3 = Red-brown lesions with some lesions containing uredinia (resistant), 4 = Red-brown lesions with most lesions containing uredinia (moderately resistant), and 5 = Tan lesions with uredinia (susceptible). We then input the matrix data into the R Pheatmap v1.0.12 package (https://cran.r-project.org/web/packages/pheatmap/index.html) to generate a heatmap and perform hierarchical clustering, enabling us to visualize the virulence patterns of *P. pachyrhizi* isolates.

## RESULTS

### Genome-wide SNP identification in *P. pachyrhizi* panel

We re-sequenced the whole-genomes of 42 diverse isolates of *P. pachyrhizi* collected throughout a wide geographic range, including Africa, Asia, Australia, South America, and the USA. The average genome coverage was 88.99% at 15X across 42 isolates, with a maximum coverage of 95.68% for isolate Pp10 (S1 Table). To investigate whole-genomic diversity among these 42 isolates, we identified SNPs by mapping the short-read sequences of each isolate along the UFV02 v2.1 reference genome assembly. The percent of mapped reads ranged from 75.53% (isolate CMES1958) to 98.48% (isolate TW72-1) (S1 Table).

To further explore the genomic context within *P. pachyrhizi*, we also included genetic variants discovered on the high-quality genome assemblies of the three isolates, K8108, MT2006, and UFV02, which were sequenced in a previous investigation [20]. In total, our study comprised 45 *P. pachyrhizi* isolates.

After applying quality control filters and minimum allele frequency (MAF ≥ 5%) (see Material and Methods section), our pipeline detected a total of 452,853 whole-genomic SNPs among 45 *P. pachyrhizi* isolates. The SnpEff software revealed the potential functional effects of these SNPs on the *P. pachyrhizi* genome (S5 Table). The majority of identified SNPs displayed either modifier or moderate impacts, with only 0.04% of variants classified as having a high functional impact. The largest proportion of SNPs was found in intergenic regions, while coding regions, such as exons, exhibited a relatively low number of SNPs.

### Molecular evidence reveals two major evolutionary lineages in *P. pachyrhizi*

An ML phylogenetic tree, using all whole-genomic SNPs, inferred phylogenetic relationships among 45 *P. pachyrhizi* isolates (Fig 1A). The topology of the phylogenetic tree showed two highly-divergent clades, each with high support bootstrap values (100). Here, we refer to these two clades as lineages Pp1 and Pp2, respectively. Notably, the two lineages were temporally separated. While lineage Pp1 comprised isolates sampled between 1972 and 1994, lineage Pp2 represented a large group of isolates collected more recently, from 2003 onwards.

Lineage Pp1 comprised four isolates: TW72-1, IN73-1, AU79-1, and HW94-1 (Fig 1A). These isolates were exclusively found in specific geographic locations, namely Asia (Taiwain and India), Australia, and Hawaii (Fig 2). Within the phylogenetic tree (Fig 1A), short branch lengths characterized lineage Pp2, with a total of 41 isolates. Showing a widespread distribution, members of the lineage Pp2 were present across Africa, Asia, South America, and the USA. (Fig 2).

**Fig 2.**
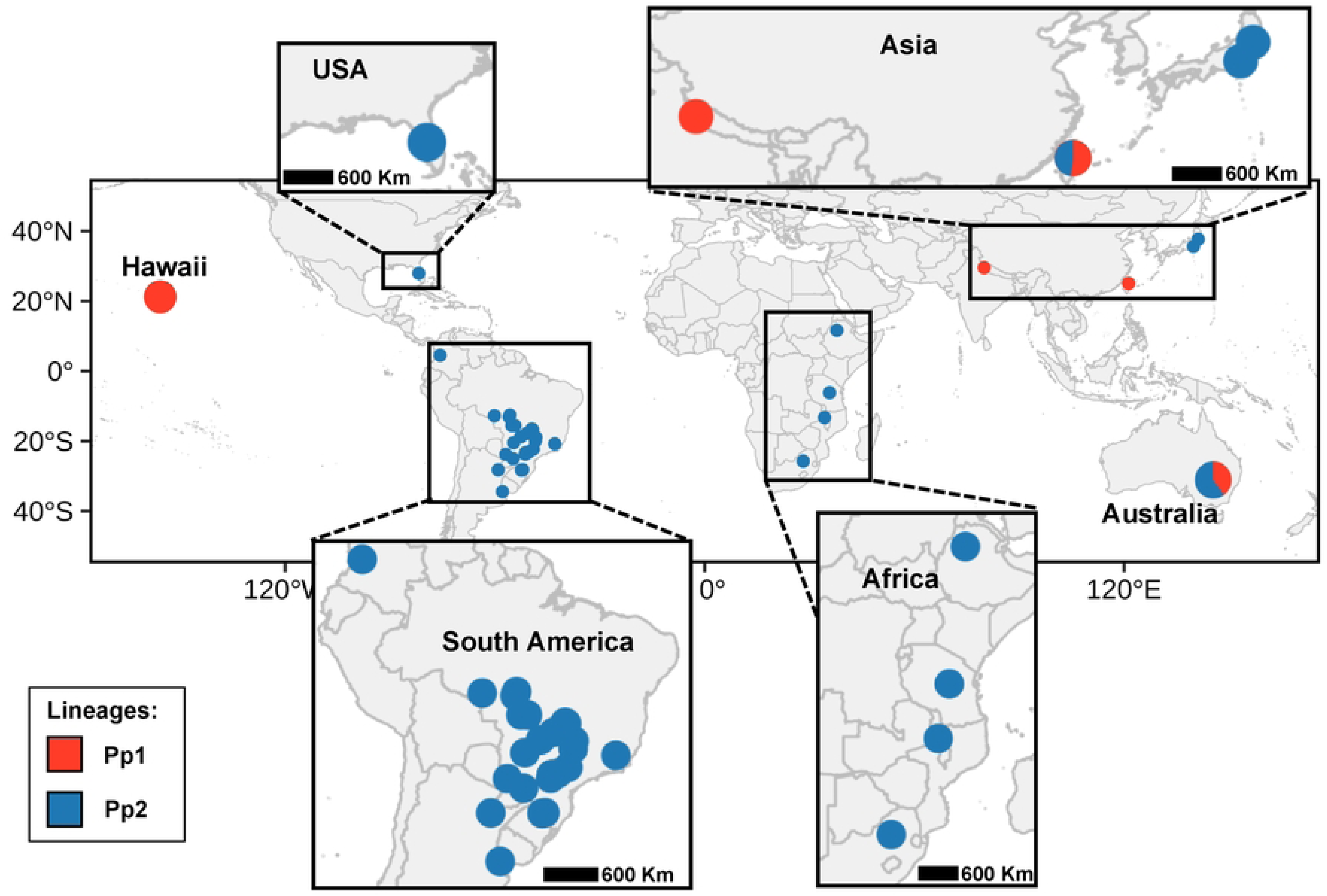
Geographic distribution of two evolutionary lineages uncovered in *Phakopsora pachyrhizi.* Color-codes: red (lineage Pp1) and blue (lineage Pp2). Each pie diagram represents a single isolate. Pie diagrams containing both colors (red and blue) represent isolates that exhibited genetic admixture levels according to fastSTRUCTURE analysis shown in Fig. 1B.

Consistent with the phylogenetic results, the FastStructure choose K.py algorithm clustered all 45 *P. pachyrhizi* isolates into two Bayesian groups (optimal K = 2; Fig 1B). As a whole, the Bayesian clustering method highly supported the presence of two divergent lineages. Importantly, the geographic origin of the isolates did not influence the placement of each Bayesian group. For example, isolates from different areas (Asia, Africa, South America, and the USA) shared similar ancestry within a single Bayesian group. Interestingly, two isolates (TW72-1 and AU79-1) contained moderate proportions of membership assigned to both Bayesian groups, indicating their genomes display certain levels of genetic admixture (Fig 1B). We also produced a PCA to illustrate genomic relationships among the 45 isolates (Fig 1C). As expected, the PCA plot revealed that the isolates form two distinct clusters, each corresponding to a described lineage.

As lineage Pp2 was the most populous clade discovered in our study, we decided to further investigate its population structure by conducting three additional analyses focusing on 41 isolates. Collectively, the three analyses (i.e. FastStructure Bayesian clustering, PCA, and Minimum spanning network) indicated a lack of genetic structure within lineage Pp2, suggesting high levels of clonal reproduction and long-distance dispersal of spores (S1 Fig).

### Genetic diversity and evidence of selection in two *P. pachyrhizi* lineages

We detected distinct patterns of molecular genetic diversity shaping the evolution of *P. pachyrhizi* lineages (Figs 3A-F and S6A-B Tables). Based on whole-genomic SNP data, lineage Pp2 exhibited significant (p < 2.2 x 10^-16^) lower levels of nucleotide diversity (mean π = 0.63 x 10^-4^) and Watterson’s Theta (mean θw = 0.29 x 10^-4^) when compared to lineage Pp1 (Figs 3A-B). This trend was also observed in both predicted effector and non-effector genes, in which lineage Pp2 showed substantially lower genetic diversity (Figs 3C-F). We found a high value of genetic differentiation between lineages Pp1 and Pp2 (Mean F_ST_ = 0.53 and Weight F_ST_ = 0.55).

**Fig 3.**
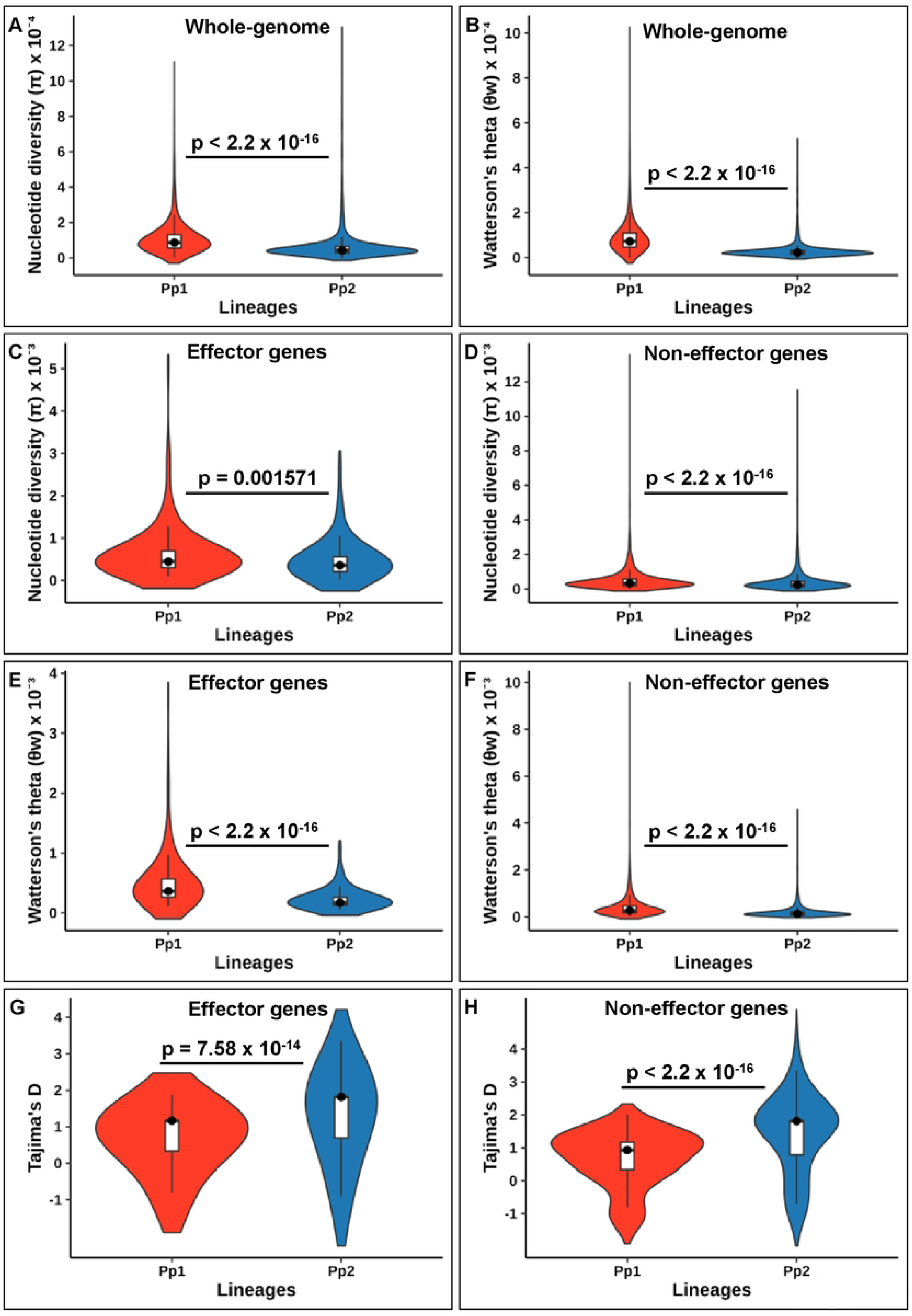
Molecular genetic diversity and Tajima’s D values for two evolutionary lineages of *Phakopsora pachyrhizi.* The analyses involved SNPs localized on three distinct regions – whole-genome, effector gene, and non-effector gene – using the UFV02 isolate as a reference genome. **A-B.** Violin plots showing the distribution of nucleotide diversity (π) and Watterson’s theta (θw) across the whole­ genome. **C-H.** Violin plots showing the distribution of π, θw, and Tajima’s D values for predicted effector and non-effector genes. Box plots were drawn inside the violin plots, with back circles representing the median. Statistical differences between the two lineages were based on the Mann-Whitney-Wilcoxon test (p-values are indicated).

Notably, Tajima’s D neutrality test suggested that *P. pachyrhizi* lineages may have evolved under different effects of natural selection (Figs 3G-H and S6A-B Tables). Although both *P. pachyrhizi* lineages exhibited positive mean Tajima’s D values within predicted effector (Fig 3G) and non-effector genes (Fig 3H), lineage Pp2 showed a significantly higher proportion of positive Tajima’s D values, indicating balancing selection.

### Genomic characterization of mating-type (MAT) loci in *P. pachyrhizi*

We used the assembled genomes of three *P. pachyrhizi* isolates K8108, MT2006, and UFV02 ([20]) to identify *MAT* genes. The three genomes together contained a total of 35 putative *MAT* genes, including pheromone receptors (*STE3.2*), pheromone precursors (*MFA*), and homeodomain transcription factors (*HD*) (S7 Table). The *STE3.2* pheromone receptor genes from *P. pachyrhizi* shared a canonical G protein-coupled receptor domain (PF02076). The repertoire of *STE3.2* gene-encoding pheromone receptors ranged from four copies (isolate K8108) to seven (isolate UFV02), while *MFA* genes were identified with only two copies per isolate. On the other hand, each *P. pachyrhizi* genome harbored two pairs of genes that encoded homeodomain transcription factors (namely HD1 and HD2), each containing a canonical homeobox KN domain (PF05920).

We constructed a schematic diagram to better illustrate the predicted structure of MAT genes in the *P. pachyrhizi* UFV02 reference genome (Fig 4A-B). The *MAT* genes of the UFV02 genome were located in physically unlinked loci designated *P/R* and *HD*. These loci were present in both the primary (Fig 4A) and secondary (Fig 4B) haplotypes of the *P. pachyrhizi* UFV02 genome. The *P/R* loci carried *STE3.2* genes-encoding pheromone receptors and their precursors (*MFA*). In contrast, the *HD* loci contained only *HD1* and *HD2* genes-encoding homeodomain transcription factors. Within the UFV02 genome assembly, HD1 and HD2 genes were connected by short DNA sequences: 776 bp in the primary haplotype and 84 bp in the secondary haplotype (Fig 4A).

**Fig 4.**
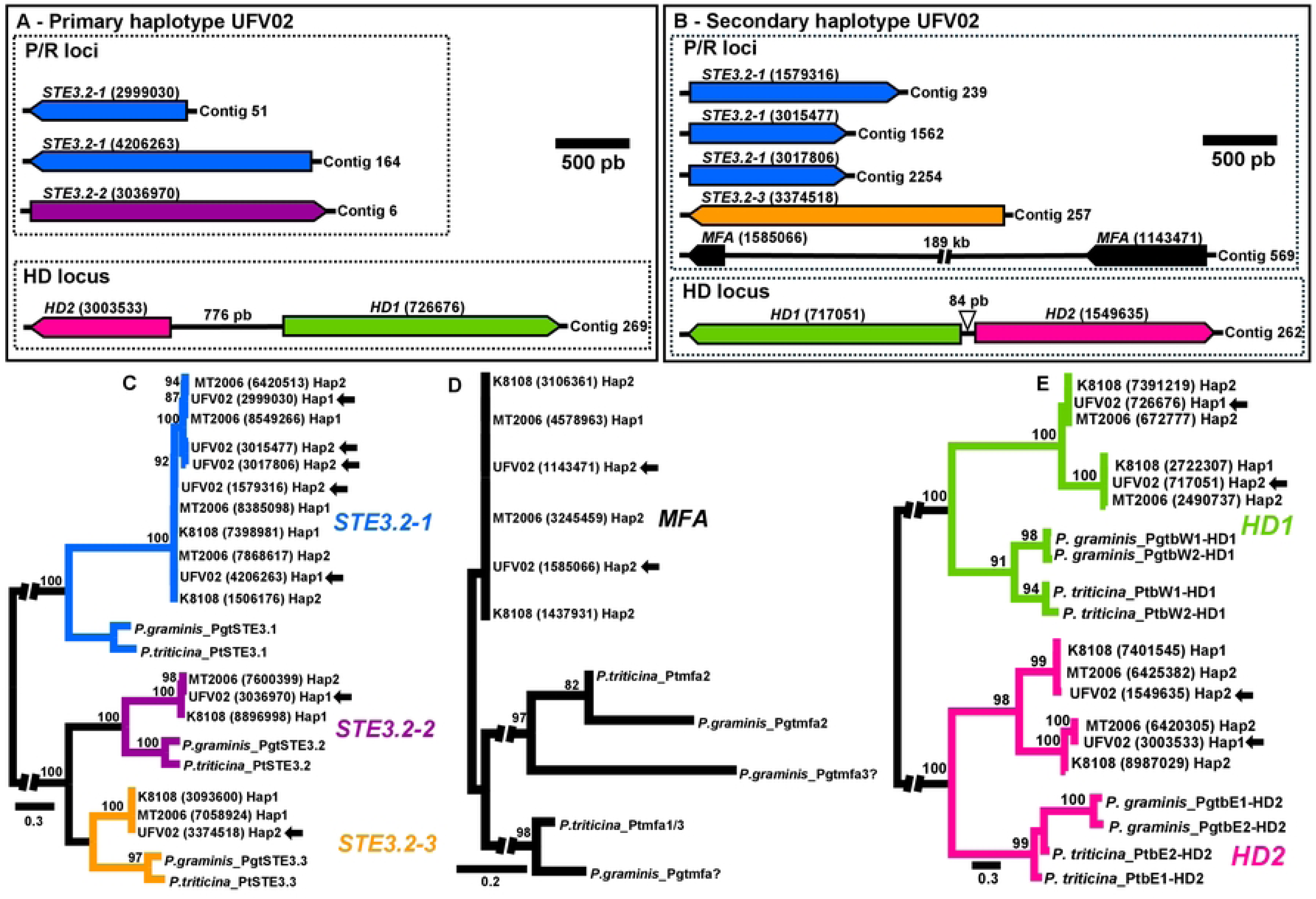
Mating-type (MAT) loci in *Phakopsora pachyrhizi.* **A-B.** Schematic diagram showing the structure of *MAT* loci on the primary and secondary haplotypes of the UFV02 reference genome. The gene lengths were drawn proportional to the scale bar indicating the number of nucleotides in base pairs (pb). **C-E.** Maximum likelihood phylogenies (unrooted tree) for three sets of *MAT* proteins in *P. pachyrhizi* isolates (UFV02, K8108, and MT2006), with *Puccinia* species as outgroups. **C.** *STE3* pheromone receptor. **D.** Pheromone precursors *(MFA).* **E.** Homeodomain (*HD*). Branch lengths are drawn to the scale bar that corresponds to the expected number of substitutions per site. Bootstrap support values are displayed above the branches (when > 80). Protein IDs are indicated in parentheses. Arrows represent the placements of *MAT* genes for the UFV02 isolate.

Our ML phylogenetic reconstructions elucidated the evolutionary relationships for each set of MAT protein sequences from *P. pachyrhizi* isolates (K8108, MT2006, and UFV02), with two *Puccinia* species serving as outgroups (Fig 4C-E). The STE3.2 pheromone receptors from both *P. pachyrhizi* and *Puccinia* species were clustered into two major highly-supported clades, with bootstrap values of 100. (Fig 4C). In agreement with previous findings [24], one such clade was named STE3.2.1, while the other was further divided into two sub-clades, STE3.2.2 and STE3.2.3. With a low bootstrap value (>80), the MAF proteins from *P. pachyrhizi* constituted a single clade, (Fig 4D). We found that HD proteins from *P. pachyrhizi* were closely related to those from *Puccinia* species, forming two main clades, each representing a homologous gene (*HD1* and *HD2*) (Fig 4E).

To gain new insights into the expression patterns of *MAT* genes during the interaction between *P. pachyrhizi* and soybean plants, we analyzed publicly accessible transcriptome data for the three isolates K8108, MT2006, and UFV02 ([20]). The expression of *MAT* genes was evaluated throughout various time points, including *in-vitro* stages (spore germination) and *in-plant* stages (soybean infection process). Overall, *MAT* genes in those three isolates exhibited varying levels of expression over the examined timeframe (S2 Fig and S8 Table). We detected a notable trend for *MAT* genes showing increasing expression levels from 72 hours post-spore inoculation. This finding was most obvious in isolate UFV02 (S2C Fig). The expression of several *STE3.2* pheromone receptors and *HD* genes were significantly up-regulated in the late stages of soybean infection, namely during sporulation.

Finally, we examined polymorphisms among the predicted *MAT* genes across whole-genome data from all 45 *P. pachyrhizi* isolates (Dataset A). Our analysis identified six SNPs within *STE3.2.1* pheromone receptor genes, while no SNPs were observed in the remaining *MAT* genes (S9 Table). By applying SnpEff software, we found that four out of six SNPs are located within intron regions of *STE3.2.1* genes, while the other two SNPs were predicted to be synonymous or missense variants.

### Virulence profiles of *P. pachyrhizi*

Our phenotyping assays identified distinct virulence patterns among four *P. pachyrhizi* isolates (AU79-1, HW94-1, IN73-1, and TW72-1) from lineage Pp1 and three isolates (CO04-2, MT2006, and UFV02) from lineage Pp2 (Fig 5, S3 Fig, and S10A-B Tables). The virulence traits of *P. pachyrhizi* isolates were analyzed on eight differential soybean PI lines, plus a susceptible soybean accession (BRS 184 or Wm82) utilized as a positive control. Both isolates, HW94-1 and IN73-1, from lineage Pp1 exhibited low profiles of virulence, with all eight soybean PI lines demonstrating immune reactions or some resistance response to these isolates (Fig 5). On the contrary, the highest levels of virulence were present in the isolate UFV02, a member of the lineage Pp2. The isolate UFV02 caused susceptible reactions in eight soybean lines, although PI 200487 (*Rpp5*) was moderately resistant to it. In terms of virulence levels, the hierarchical clustering dendrogram (Fig 5) did not reflect the phylogenetic placements of the isolates as shown in Fig 1A

**Fig 5.**
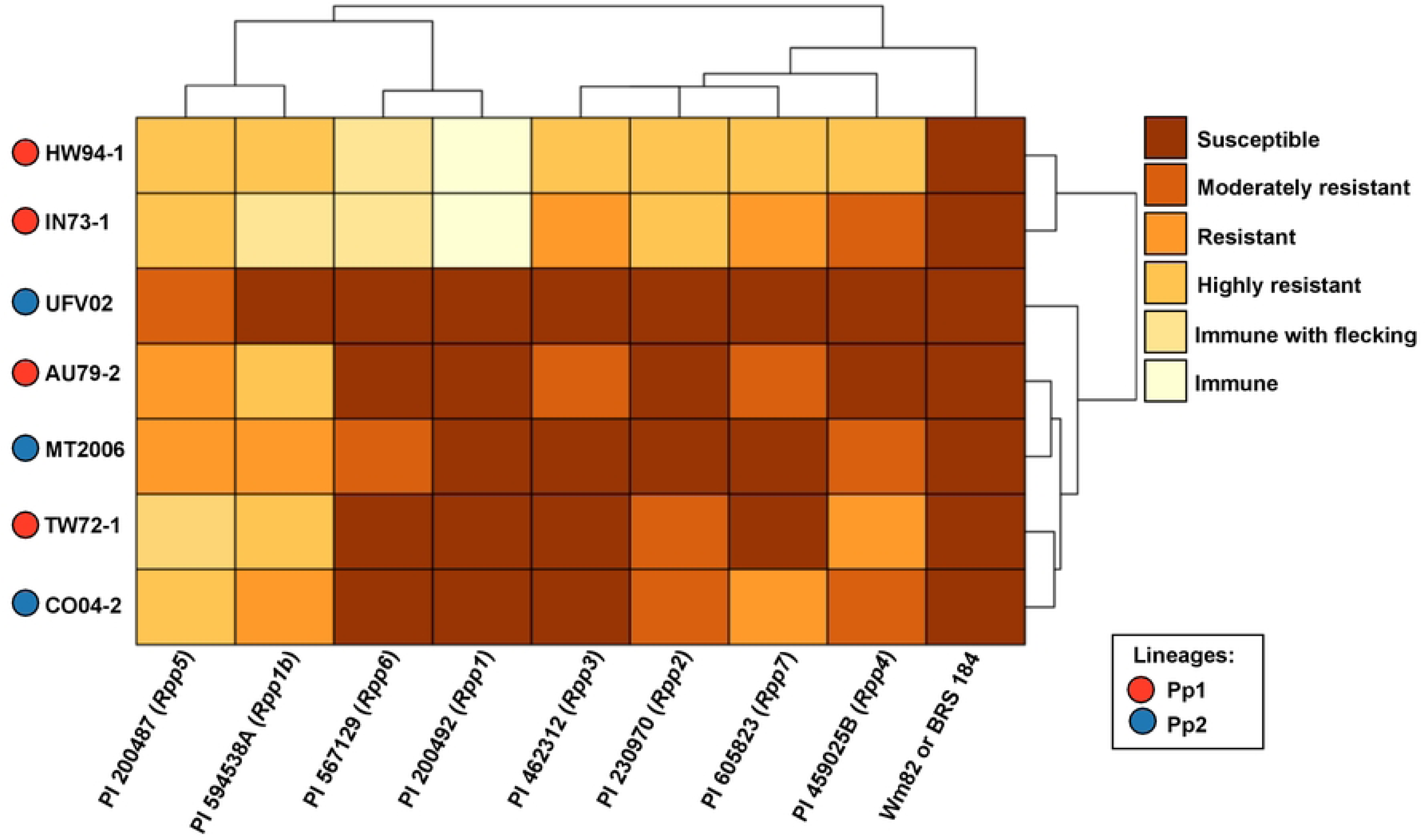
Heatmap illustrating the virulence levels of *Phakopsora pachyrhizi* isolates from two major lineages (Pp1 and Pp2) on differential soybean genotypes. The x-axis shows eight soybean Pl accessions with distinct *Rpp* genes: **Pl** 200492 *(Rpp1),* Pl 594538A *(Rpp1b),* Pl 230970 *(Rpp2),* Pl 462312 *(Rpp3),* Pl 4590258 *(Rpp4),* Pl 200487 *(Rpp 5),* Pl 567129 *(Rpp6),* and Pl 605823 *(Rpp7)*. A susceptible genotype (BRS 184 or Wm82) was included as a positive control. The y-axis represents six *P. pachyrhizi* isolates: AU79-1, C004-2, HW94-1, IN73-1, MT2006, TW72-1, and UFV02. Color intensity reflects the virulence level of each *P. pachyrhizi* isolate on the respective soybean genotype. Dendrograms are included to illustrate the hierarchical clustering of soybean genotypes (columns) and *P. pachyrhizi* isolates(rows).

## DISCUSSION

### Two evolutionary lineages drove the diversification of *P. pachyrhizi*

Previous studies of genetic diversity in *P. pachyrhizi* have relied on microsatellite markers [17], AFLPs [18], as well as specific genomic regions such as ITS [16, 15]. This study represents a pioneering effort to acquire whole-genome re- sequencing data from multiple *P. pachyrhizi* isolates. Our results provide a comprehensive view of the genetic diversity in *P. pachyrhizi* at the genome-wide level.

In our study, phylogenetic reconstructions, based on whole-genomic SNP data, uncovered two divergent evolutionary lineages of *P. pachyrhizi* distributed across diverse global regions. It is well-established that populations of plant-pathogenic fungi undergo rapid evolutionary changes, often leading to the emergence of locally adapted lineages in response to selection pressure from host species [50]. Indeed, the evolution of virulent clonal lineages has been documented in invasive rust fungal populations, including *Puccinia graminis* f. sp*. tritici* [51], *Uromyces transversalis* [52], and *Puccinia triticina* [53]. Our findings align with previous studies that identified genetically distinct clusters in *P. pachyrhizi* [17, 19]. Utilizing both ITS and ARF sequences, molecular analyses showed that *P. pachyrhizi* genotypes from four geographic regions — Africa, America, Asia, and Australia — formed distinct clades that were not influenced by geographic locations of the *P. pachyrhizi* isolates [19].

The long-distance transport of airborne spores via air currents has been a key dispersal mechanism for *P. pachyrhizi* [5, 14). This phenomenon has potentially contributed to the widespread distribution of members of the two evolutionary lineages (Pp1 and Pp2) in *P. pachyrhizi* across large geographic distances. Notably, lineage Pp1 was predominantly dispersed in the Eastern Hemisphere, with its members established in countries such as Australia, India, and Taiwan. Long-distance dispersal events likely explain the presence of one member of lineage Pp1, observed in the Hawaii archipelago. With a larger geographic distribution, lineage Pp2 exhibits a remarkable ability to disseminate fungal spores across vast distances. Through airborne inoculum dispersal, members of the lineage Pp2 may have reached major soybean-producing centers in Africa, South America, and the continental USA. Consistent with a general trend observed in prior works [15, 18], we found a lack of genetic structure across broad geographic ranges within each of the evolutionary lineages, Pp1 and Pp2, in *P. pachyrhizi*. This absence of spatial genetic structure in *P. pachyrhizi* lineages is likely due to the high propagation rates of clonal spores among soybean-growing areas located on large geographic scales. The maintenance of *P. pachyrhizi* clonal lineages in fields may be facilitated by alternative host species, which serve as inoculum reservoirs during planting seasons [2]

Despite their close phylogenetic relationships, limited gene flow was observed between the two *P. pachyrhizi* lineages, Pp1 and Pp2, as indicated by high F_ST_ values. Within lineage Pp2, low levels of polymorphism were detected, with only a few SNPs distinguishing different isolates. The reduced genetic diversity suggests that lineage Pp2 displays highly clonal reproduction. In contrast, a significant accumulation of genomic polymorphism was observed among isolates in lineage Pp1. The high levels of genomic variability within lineage Pp1 may be attributed to its presence in regions with low-intensity soybean production, where relaxed selective constraints could have promoted increased genetic variation. The genetic diversity of *P. pachyrhizi* may also reflect mechanisms such as somatic recombination via hyphal anastomosis [54] and intense activity of transposable elements in the dikaryotic urediniospores [20].

Temporal shifts are thought to drive the emergence of new genotypes and the selection of specific gene loci to elevate virulence in plant-pathogen populations [55]. The temporal divergence between the two lineages Pp1 and Pp2 in *P. pachyrhizi* is an interesting discovery. Our findings indicate that lineage Pp1 cluster *P. pachyrhizi* isolates obtained from several decades ago (1972 -1994), while lineage Pp2 includes more recently collected isolates (2003-2017). We propose two possible scenarios to explain the temporal separation observed in the two *P. pachyrhizi* lineages. The first scenario suggests that our sampling efforts may have been insufficient to catch recent isolates of *P. pachyrhizi* belonging to lineage Pp1. This scenario implies that lineage Pp1 continues to circulate in the current soybean-producing zones. On the other hand, the second scenario postulates that lineage Pp1 may represent an ancestral group, and its parental members could have evolved in response to host interactions over time, subsequently giving rise to lineage Pp2, which is now prevalent across the world. The impact of temporal differentiation on the population structure of plant-pathogens has been reported. For example, in the crown rust fungus (*Puccinia coronata* f. sp*. avenae*), populations sampled in the USA during 1990 and 2015 formed two distinct genetic clusters, each comprising isolates from a single collection year [55]. In this case, the latter *P. coronata* population (2015) likely derived from the 1990 population and acquired a substantial increase in virulence levels [55].

Previously, evidence indicated that *P. pachyrhizi* appears to be originated in Asia [5], with its earliest records from 1902 in Japan [9]. We speculate that lineages Pp1 and Pp2 may have initially emerged in Asia and later spread to geographic regions worldwide. The co-existence of both evolutionary lineages Pp1 and Pp2 in Asia implies that their evolutionary histories may have been interconnected over ancient periods in that region. However, extensive sampling of additional *P. pachyrhizi* isolates, especially in the Asian continent, is required to determine the precise geographic origin for lineages Pp1 and Pp2 of *P. pachyrhizi*.

### Genetic admixture among *P. pachyrhizi* lineages

The origin of two *P. pachyrhizi* isolates, TW72-1 (from Taiwan) and AU79-1 (from Australia), is likely related to genomic admixture involving the lineages Pp1 and Pp2. The fastStructure analysis revealed that TW72-1 and AU79-1 share ancestral origins to both lineages Pp1 and Pp2 (Fig 1B). Genetic admixture in rust fungi may be a consequence of multiple mechanisms, including sexual recombination and somatic hybridization [56]. A recent study utilizing fully nuclear-phased genome assemblies demonstrated that somatic hybridization through nuclear exchange led to the emergence of wheat leaf rust (*Puccinia triticina*) lineages [57]. The mechanisms involving the genetic admixture of TW72-1 and AU79-1 are not fully explained in our study. However, one possible explanation is that members of lineages Pp1 and Pp2 may be mating-compatible partners and can outcross, producing new recombinant genotypes (e.g. TW72-1 and AU79-1). This hypothesis would be supported by the presence of different *MAT* alleles among isolates of lineages Pp1 and Pp2, although our analysis revealed reduced allelic variation within *MAT* genes of *P. pachyrhizi* lineages (S9 Table). Specifically, only the *STE3.2.1* pheromone receptor genes showed some polymorphisms, while other *MAT* genes exhibited no allelic variations among isolates in lineages Pp1 and Pp2. Therefore, sexually compatible outcrossing between lineages Pp1 and Pp2 appears a less likely hypothesis to elucidate the origin of TW72-1 and AU79-1. Alternatively, somatic hybridization through nuclear exchanges may be a more plausible scenario to explain the genetic admixture observed in TW72-1 and AU79-1. Considering this possible scenario, isolates TW72-1 and AU79-1 may harbor dikaryotic spores containing two different nuclear haplotypes that were derived from lineages Pp1 and Pp2, independently. The recognized occurrence of hyphal anastomosis followed by nuclear migration in germ tubes of *P. pachyrhizi* spores [54] further reinforces the possibility of a somatic hybridization event among *P. pachyrhizi* lineages. Similar incidents associated with somatic hybridization via nuclear exchange have been documented in other fungi rust, for instance, *Puccinia triticina* [57], *Puccinia graminis* f. sp. *tritici* [58] and *Puccinia coronata* f. sp*. avenae* [59]. Although the sexual cycle of *P. pachyrhizi* has not been characterized, its role should not be ruled out as a factor to explain the genomic admixture observed in TW72-1 and AU79-1.

### Tetrapolar mating types in *P. pachyrhizi*

In basidiomycete fungi, the role of *MAT* genes in controlling sexual compatibility is well-established [27]. In line with previous findings in other basidiomycete fungi — *Puccinia* species [25] and myrtle rust *Austropuccinia psidii* [26] — our analyses confirmed that *P. pachyrhizi* possesses a tetrapolar mating system that likely regulates sexual compatibility. Indeed, within the phylum Basidiomycota including rust fungi, a tetrapolar mating system appears to be an ancestral trait [25]. The tetrapolar mating system of *P. pachyrhizi*, composed of physically unlinked HD and P/R loci, is present in both primary and secondary haplotypes of the genome assemblies. This finding supports a common pattern observed in rust fungi [26, 25]. Typically, in the cereal rust fungi genomes, *MFA* pheromone genes are found close to their respective *STE3.2* gene-encoding receptors on the chromosomes [25]) On the contrary, the MFA genes of *P. pachyrhizi* were located on distinct contigs separated from the *STE3.2* receptor genes. This unexpected finding likely reflects fragmented genome assemblies of *P. pachyrhizi*, which may have prevented us from achieving the full understanding of the genomic architecture of *MAT* genes in this species. While two *MFA* pheromone genes were identified in each examined *P. pachyrhizi* isolate, it is possible that additional *MFA* copies are present within their genomes. The short length of *MFA* proteins (80 amino acids) in *P. pachyrhizi* further complicates their identification through bioinformatic tools like BlastP.

The predicted *MAT* genes in *P. pachyrhizi* are potentially functional, as evidenced by their significant expression levels observed in the transcriptome data. The high expression of MAT genes during the later stages of soybean*-P. pachyrhizi* infection is consistent with previous observations in the poplar leaf rust fungus *Melampsora larici-populina* [60] and wheat rust fungi, e.g. *Puccinia* species [25]. The notable expression patterns of *P. pachyrhizi* MAT genes raise interesting questions about how *MAT* genes might contribute to urediniospores formation over the late stages of the *P. pachyrhizi* infection cycle on soybean. A recent study, focusing on rust fungi within the genus *Puccinia*, suggested that *MAT* genes may play a crucial role in regulating the pairing of two haploid nuclei during the development of new dikaryotic urediniospores in the late infection phases [25]. If *MAT* genes are indeed involved in the formation of *P. pachyrhizi* asexual urediniospores, experimental analyses are needed to confirm their functional roles.

### Virulence profiles of *P. pachyrhizi*: implications for disease management

Experiment-based evidence has revealed the existence of several potential pathogenic races of *P. pachyrhizi*, with varying levels of virulence [6, 7]. Our findings demonstrated that different isolates from lineages Pp1 and Pp2 are virulent in most differential soybean lines, indicating their notable adaptive ability to overcome resistance conferred by *Rpp* genes. The large genome size of *P. pachyrhizi*, coupled with a high abundance of transposable elements, is thought to play a key role in the adaptability and virulence of the pathogen [20].

The emergence of new *P. pachyrhizi* lineages demand the attention and surveillance of phytopathologists to update ASR-resistance breeding or crop protection approaches. To control ASR, the primary strategies have been fungicide applications and the use of resistant soybean varieties [8]. The effective monitoring and managing of ASR is indispensable since *P. pachyrhizi* populations may rapidly adapt to local conditions, manipulating host defense mechanisms and giving rise to virulent strains that can quickly spread to neighboring regions via wind-dispersed spores. When screening soybean germplasm for resistance to ASR, breeding programs should consider the genomic variation among *P. pachyrhizi* isolates within the lineages Pp1 and Pp2 as described in our study.

## SUPPORTING INFORMATION

**S1 Fig. Population structure for 41 *Phakopsora pachyrhizi* isolates belonging to lineage Pp2.** The dataset consisted of 115.065 SNPs distributed across the genome. **A.** Results from fastSTRUCTURE analysis reveal a single Bayesian cluster (K=1). Each colored horizontal bar represents the ancestry proportion of an isolate to its respective Bayesian cluster. **B.** Plot of the first two principal components, with eigenvalues for each component shown in parentheses. **C.** Minimum spanning network (MSN) depicting genetic relationships among *P. pachyrhizi* isolates, based on Euclidean distance via R Poppr package.

S2 Fig. Expression patterns of mating-type (MAT) genes in three *Phakopsora pachyrhizi* isolates during soybean infection. Isolates: **A.** K8108, **B.** MT2006, **C.** UFV02. Gene expression was assessed in both *in-vitro* and *in-plant* stages. The *in-vitro* stages included spore and appressorium samples (Ap_out), while *in-plant* stages spanned 10 to 288 hours post-inoculation (HPI). The *in-plant* appressorium samples (Ap_on) were exclusively evaluated for isolate K81081. Expression levels were calculated as Log2 fold change (Log2FC). The x-axis represents distinct time points at which gene expression was evaluated, while the y-axis represents the distinct MAT genes. Color intensity reflects expression levels according to the color palette. Gene expression data were derived from RNA-seq libraries analyzed in a previous study [20].

S3 Fig. Lesion profiles of soybean genotypes 14 days after spore inoculation, with *Phakopsora pachyrhizi* isolates came from two independent lineages (Pp1 and Pp2). Phenotyping evaluations included eight soybean PI accessions carrying seven different *Rpp* loci (*Rpp*1-*Rpp*7), with a susceptible genotype (BRS 184 or Wm82) used as a control. Scale bar = 0.5 cm.

S1 Table. List of *Phakopsora pachyrhizi* isolates, with geographical origin, collection date, read filtering, number of mapped reads, and genome coverage.

S2 Table. *Puccinia* mating-type (MAT) genes used as queries in BlastP searches.

S3 Table. Differential soybean accessions used in the virulence assessment of *Phakopsora pachyrhizi* isolates.

S4 Table. Reaction categories based on quantitative responses produced by *Phakopsora pachyrhizi* isolates.

S5 Table. Predicted impact of SNPs on whole-genome data from 45 *Phakopsora pachyrhizi* isolates. The analysis was performed in SnpEff software, using UFV02 isolate as a reference genome.

**S6 Table. Genetic diversity measures for two evolutionary lineages of *Phakopsora pachyrhizi.*** The analyses were based on SNPs present in three different regions: whole-genome, effector genes, and non-effector genes. Statistical differences between the pair of lineages were assessed by the Mann-Whitney-Wilcoxon test. **S6A.** Overview of nucleotide diversity (π), Watterson’s theta (θw), and Tajima’s D values for two evolutionary lineages. **S6B.** Nucleotide diversity (π), Watterson’s theta (θw), and Tajima’s D values for predicted genes (effector and non-effector) in the UFV02 genome reference.

**S7 Table. Mating-type (MAT) genes in three Phakopsora pachyrhizi isolates (UFV02, K8108, and MT2006).**

**S8 Table. Expression profiles of mating-type (MAT) genes in three *Phakopsora pachyrhizi* isolates (UFV02, K8108, and MT2006).** Gene expression data came from RNA-seq libraries analyzed in a prior study [20]. Significant p-values (< 0.05) are indicated in green.

**S9 Table. Predicted impact of SNPs on mating-type (MAT) genes of *Phakopsora pachyrhizi*.** The analysis was conducted in SnpEff software and gene models from the UFV02 were taken a reference.

S10 Table. Infection types of soybean genotypes 14 days after spore inoculation, with *Phakopsora pachyrhizi* isolates came from two independent lineages (Pp1 and Pp2). The phenotype evaluation included eight soybean PI accessions carrying seven different *Rpp* loci (*Rpp*1-*Rpp*7), with a susceptible genotype (BRS 184 or Wm82) used as a control. **S10A.** Virulence levels of *Phakopsora pachyrhizi* isolates. **S10B.** Sporulation level (SL), Number of uredinia per lesion (NoU), and percentage of lesions with uredinia (%LU) for two *Phakopsora pachyrhizi* isolates MT2006 and UFV02. The quantitative data were obtained from 3 biological replicates (one leaf per plant), with 30 evaluated lesions per leaf, totaling 90 lesions per soybean differential genotype.

## ACKNOWLEDGEMENTS

The work (proposal: 10.46936/10.25585/60000959) conducted by the U.S. Department of Energy Joint Genome Institute (https://ror.org/04xm1d337), a DOE Office of Science User Facility, is supported by the Office of Science of the U.S. Department of Energy operated under Contract No. DE-AC02-05CH11231. The study was supported by the Brazilian Agricultural Research Corporation - EMBRAPA (proposal 03.17.03.004.00.00) and Bayer SAS/EMBRAPA (proposal 33.17.00.113.00.00). The National Council of Scientific and Technological Development - CNPq and the Brazilian Ministry of Science, Technology and Innovation - (MCTI) funded project 440782/2022-8 and supported FCMG (PQ 312214/2022-7) and VDR (DTI-A 383533/2023-6). This work was also partially funded by the USDA-ARS project 8044-22000-051-000D. SD was supported by a grant of the French Plan Investissement d’Avenir (PIA) Lab of Excellence ARBRE [ANR-11-LABX-0002- 01]. We acknowledge Dr. Glen Hartman and Dr. H. M. Murithi, who kindly provided the isolates from Africa, Argentina and the US; Dr. Y. Yamaoka for the Japanese isolates, and Dr. Sergio Herminio Brommonschenkell for the UFV02 isolate.

## Author contributions

**F.C.M.G, M.L, R.T.V, I.V.G, U.S, C.S, and S.B:** Conceptualization, Funding acquisition, Manuscript revision. **F.M.C and M.K.K:** Methodology, Isolation and cultivation of fungal isolates. **M.K.K:** Methodology, DNA extraction and quality analysis, Phenotypic characterization **K.F.P.**: Cultivation of fungal isolates, Phenotypic characterization, Manuscript revision. **I.V.G, A.L and K.B:** Methodology, Re-sequencing analysis and In-silico analysis in the JGI platform. **C.V.G and M.C.M**: Fungal spore sampling. **E.G.C F:** Methodology, Identification of genetic variants. **V.D.R**: Data curation, Formal analysis, Visualization, Writing – original draft. **F.C.M.G**: Writing – review & editing and Supervision. All authors read and approved the submitted version.

## DATA AVAILABILITY

Whole-genome re-sequencing data have been deposited at the JGI portal (https://genome.jgi.doe.gov/portal/) under project IDs available in S1 Table.

## Disclaimer

Mention of trade names or commercial products in this publication is solely for the purpose of providing specific information and does not imply recommendation or endorsement by the U.S. Department of Agriculture. The USDA is an equal opportunity provider and employer.

